# Investigation of neural functional connectivity in thick acute mouse brain slices with novel multi-region 3D neural probe arrays

**DOI:** 10.1101/2024.01.18.576320

**Authors:** Wesley Charles Smith, Zoia Naumkina, Hyo Geun Shin, Ui Kyu Chae, SeungHun Lee, Jung-Hoon Park, Yak Dol Cho, Ji Wan Woo, Seok Kyu Kwon, Soo Jin Oh, Min-Ho Nam, Tae Song Kim, Il Joo Cho

**Affiliations:** Brain Science Institute, Korea Institute of Science and Technology (KIST); Department of Convergence Medicine, College of Medicine, Korea University; Department of Biomedical Engineering, Ulsan National Institute of Science and Technology; Research Animal Resource Center, KIST

**Keywords:** *ex vivo*, *3D probes*, *functional connectivity*, *extracellular electrophysiology*, *optogenetics*

## Abstract

There are significant limitations in investigating complex neural circuits *in vivo*, including drawbacks to midline-adjacent surgeries, limited accessibility to deep brain regions and number of feasible regional targets for simultaneous recordings, and analytical or experimental biases from recording one columnar plane. On the other hand, recording extracellular neural signals *ex vivo* or *in vitro* using planar microelectrode arrays (MEAs) only permits slice surface recordings, and since conventional slices under 400 μm-thick or dissociated cultures are used, no experiments contain a physiological multi-region circuit, drastically limiting conclusions about connectivity and pharmacology. Using thick, tract-preserving acute brain slices to record otherwise unassailable neural circuits *ex vivo* combines the strengths of both types of experiments, but is assumed to precipitate ischemic injury due to oxygen scarcity within the slice. Here, we report the first application of custom, multi-region silicon neural probe arrays to record spontaneous activity & optogenetically-induced functional connectivity acrosshe mesocorticolimbic pathway within tract-preserving 800 μm sagittal mouse brain slices, compared with 400 μm slices, among three brain regions: the ventral tegmental area (VTA), ventral striatum (VS), & medial prefrontal cortex (mPFC). We show that most single-unit signals are an order of magnitude below the noise floor seen using silicon probes *in vivo*, providing unit yields far higher than previously assumed, allowing for a deep functional understanding of acute slice condition compared to the assumed deterioration due to ischemia. Overall, our method allows for acute circuit manipulations beyond what is available in vivo, with far more information than conventional slice preparations.

## Introduction

Currently, neurophysiology is viewed primarily as an *in vivo* application of increasing channel counts in silicon neuroprobe arrays and development of faster cell imaging sensors or as a rote method of ascertaining channel functionality intracellularly in *ex vivo* preparations.

Despite obvious strengths, *in vivo* electrophysiology has multiple drawbacks that are inextricably linked. Columnar arrays with hundreds of channels implicitly have biased location sampling and can’t record many functionally connected nuclei or sub-circuits due to geometric constraints, putting a limit on the number of sites or regions that can be simultaneously recorded(1, 2), especially deep targets. These techniques are paired with upper physical limitations on what can be delivered as a brain stimulus to a mouse or rat, as the investigator cannot control the entire inner milieu *in vivo*. As well, neurons are recorded with sharp electrodes, fully occluded, hence the popularity of the aforementioned imaging techniques. Subsequently, this causes bleeding and inflammation(3), which interferes with recording accuracy and subject behavior, especially when probing midline-adjacent, heavily vascularized tissue(4). Motion artefacts(5) and inflammation cause signal drift(6), high RMS background noise(7, 8), and lead to sparse sampling and amplify the seldom-discussed “dark neuron” problem(9), and skew analysis towards experimenter-determined ’robust’ neurons.

Attempts to circumvent some of these issues through activity imaging will still preclude recording vast or deep circuit targets, and will have fundamentally slower timescales and require more artifice to catch up to electrophysiology, while not even supplying basic data like LFPs(5, 10). The sparse sampling problem is worsened by dealing with low resolution color rather than voltages, while for equivalent epochs, raw data can be several times larger, exacerbated by ultra-fast cameras and voltage imaging (11). Finally, an often ignored requisite is the implied use of transgenic animals or light excitable indicator chemicals which undoubtedly have physiological consequences(12) but are not often the subject of investigation, and eventually become lurking variables and side effects.

At the other end of the physiology spectrum, *ex vivo* slice and dissociated culture recordings cannot recapitulate a physiological circuit at all, and although organotypic slice preps may use multielectrode arrays (MEAs), only the surface touches the recording channels and all tracts are severed. Most *ex vivo* experiments are very granular, missing the forest for the trees in assumptions about pharmacological effects with a focus on channel function, without considering circuit or region function. There is also selection bias towards healthy neurons(13) and a large dependency on experimenter skill and knowledge in preparing the entire environment for the tissue including: extraction, solution preparation, temperature sensitivity, and handling, which has a steep learning curve and readily yields different experimental outcomes. As such, some experimenters can intracellularly record a handful of cells at once(14), while most will never master the technique in such a deep way. And, despite meticulous selection of healthy neurons, their sampling is concentrated near the surface of the slice, where tissue damage from dissection and sectioning is evident, also exposing healthier tissue to leaking cytokines, signaling molecules, and other ectopic cytotoxic debris (15).

We propose a protocol and platform for using tract-preserving 800 µm thick acute sagittal brain slices to record a normally unassailable deep brain circuit, the mesocorticolimbic path and its hubs the VTA, mPFC, and VS (16), and using optogenetic stimulation of ChR2 under the Thy1 promoter (17), compare functional connectivity to that of a 400 µm slice. This combines the strengths of a more intact *in vivo* prep’s silicon probe analysis methods with the experimental freedom of slice and our custom neuroprobe arrays to demonstrate the applicability of this technique to all manner of other circuits. It’s unknown how efficacious this is, considering that most slice methods use 400 µm or thinner slices due to cell visualization issues and concerns about ischemic injury due to hypoxia deep in the slice (18–20), and there is insufficient electrophysiological data on thick slice viability(21). This insufficiency yields wildly differing approaches to quantify tissue oxygenation and sources of ischemia (22, 23) while ignoring the actual issue of physiology, hence, creating a blind-spot in the literature.

We demonstrate that the VTA to VS and mPFC portion of the mesocorticolimbic circuit in the MFB (medial forebrain bundle) retains a greater proportion of optically responsive units in the 800 µm thick slice only, while we also see *in vivo*-like spontaneous activity generally, with little evidence of degraded activity or hypoxia. Interestingly, the mPFC to VS and VTA optogenetic stimulation path has equivalent proportion of responsive units in both slice thicknesses. The combined strength of intact circuits, multiple regions recorded with custom arrays, experimental on-the-fly optical manipulation, multiple perfusable drugs, and ACSF manipulation with the ability to correlate any of these with prior behavior and pharmacological tests enable this to take the best of *in vivo* and *ex vivo* pipelines and push forward novel ways of approaching drug testing and neural decoding methods at the essential circuit level.

## Materials and methods

### Rationale, fabrication, construction, and electroplating of the 3D neuroprobe array

As the goal of the project was to successfully demonstrate multi-region, circuit- preserving, deep-target *ex vivo* slice electrophysiology that would normally be impossible to record with conventional commercial methods while using silicon probes, we delineated that a single hemisphere’s mesocorticolimbic pathway would be a recording target that would both satisfy those criteria while having all its hub regions within two types of slice thicknesses, albeit at different levels of integrity. This sagittal slice that spans the entire rostro-caudal brain preserves a greater degree of the MFB (mfb on Fig. 1a, Fig. 1b) only in the 800 µm slice condition (Fig. 1c, large brackets) as compared to the 400 µm slice condition (Fig. 1c, small brackets), and if started at 100 µm past midline, contains the VTA (Fig. 1c, bottom), mPFC (Fig. 1c, top), and VS (Fig. 1c, middle), all hubs of this pathway. In this manuscript, these hubs and their associated traces will generally be color-coded as: blue/mpFC, red/VS, and black/VTA for ease of communication. This novel preparation also requires a new design of silicon probe to record from; therefore we present the following methods to construct such a neuroprobe. By demonstration of this method and result for one circuit, its strength and validity may further be shown in other pathways for various paradigms and neuroscientific inquiries.

**Figure 1.**
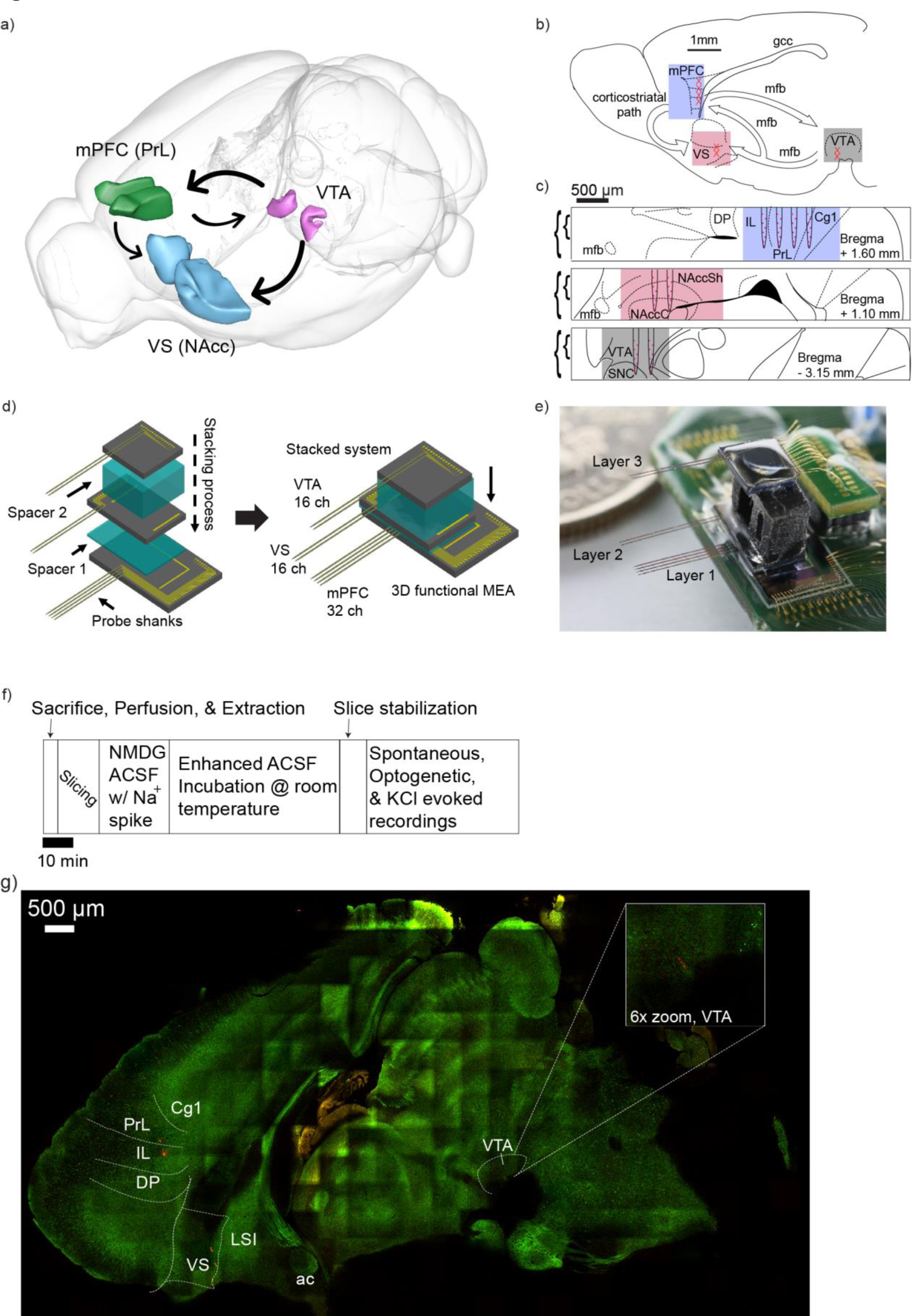
*Slice, probe assembly, and experimental schematics.* (a) Scalable Brain Atlas schematic of the principle regions recorded; VTA = ventral tegmental area, mPFC = medial prefrontal cortex, PrL = prelimbic cortex, VS = ventral striatum, NAcc = nucleus accumbens. (b), Sagittal slice schematic with color-coded regions for ease of association and simplified medial forebrain bundle (mfb) diagram. (c) Coronal sections of mPFC (top), VS (middle) and VTA (bottom) regions within the same sagittal slice, with slice extent indicated at brackets for 400 µm (inner, small brackets) and 800 µm (outer, large brackets). Note the large channel spacing on each set of probes. DP = dorsal peduncular cortex, IL = infralimbic cortex, Cg1 = cingulate cortex area 1, NaccC = nucleus accumbens core, NAccSh = nucleus accumbens shell, SNC = substantia nigra pars compacta. (d) 3D probe stacking process with spacers in blue. (e) Complete probe with extra PCB for wirebonding on the far right. (f) Recording day timeline. (g) Multiphoton image of the slice recorded in Figures 2-4. Thy1-ChR2 expressing cells in green, DiI probe tracks in red.

The fabrication of a 3D neural probe array is divided mainly into three steps: (1) the fabrication of three planar 2D neural probe MEAs following methodology as previously published by our group (24), (2) the assembly of the 3D MEA by stacking the 2D MEAs with specially printed spacers(1), and (3) the formation of electrical connections. First, to fabricate the multifunctional MEMS neural probe, we used a four-inch silicon-on-insulator (SOI) wafer with 40 μm thick top silicon. Next, a 400 nm thick electrical insulation layer (SiO2) was deposited through plasma-enhanced chemical vapor deposition. After consecutive Ti/Au (200 Å/3000 Å) deposition and patterning of the signal line, a second 400 nm thick SiO2 layer was deposited on top. After patterning the wide-spaced electrode sites (Fig. 1b, all insets), the insulation layer on the electrode sites was etched using reactive ion etching (RIE), and Ti/Pt (200 Å/1500 Å) was deposited on the electrode sites through a lift-off process. After patterning of the top Si layer in the shape of a neural probe, the structure was released through consecutive RIE and deep RIE processes.

Second, the 3D system was formed by stacking and bonding the 2D independent probes. One of the features of the system is the specificity and accuracy of the alignment of each probe above the others. For this purpose, a special aligner was designed to control orientation. The equalizer structure was positioned on the side of the probe and supported the entire system and after assembly was completed, it was easily removed. The first probe body was fixed to a PCB (Fig. 1d, left). Then, the bottom spacer was settled and the second neural probe was fixed on top of the spacer. Likewise, the same process was performed for the second spacer and third neural probe to complete the array (Fig. 1d, right).

Finally, the electrical pads on the body of the bottom neural probe were directly wire-bonded to pads on the custom PCB (iBond5000, MPP Tools), and two upper neural matrixes were connected to the main PCB through two-step bonding technology (Fig. 1e). 64ch board-to-board connectors (Omnetics) were soldered onto the 3D neural platform PCB to enable repeated reattachment and reuse with Intan Headstages (64ch C3315, Intan Technologies) and their output with SPI cables (Intan Technologies).

To evaluate the performance of the 3D neuroprobe array, we measured the impedance of the microelectrodes with a frequency sweep (10 Hz-10 kHz) using an impedance analysis system (nanoZ, Neuralynx, Bozeman, Montana, USA). We electroplated Pt black on the Pt microelectrodes to improve the quality of their recording. The Pt black plating solution contained 3% hexachloroplatinic acid hydrate (520896-5G, Sigma-Aldrich, USA), 0.025% HCl (4090-4400, DAEJUNG, South Korea), and 0.025% lead acetate (316512-5G, Sigma-Aldrich, USA) in deionized water. The tips of the neural probe system were immersed in the plating solution with a reference electrode (Ag/AgCl wire) and a counter electrode (Pt wire). The Pt microelectrodes were electroplated selectively by applying the electrical potential (–0.2 V, 35 s) through a potentiostat (PalmSens3, PalmSens, Netherlands) to reach a mean impedance of 41.15±12.02 kΩ (standard deviation indicated). Prior to recordings, probes were coated with fluorescent dye DiD (Invitrogen); after recordings they were cleaned with Tergazyme/ddH2O (Sigma) for re-use.

### Solution preparation

Three main solutions were prepared: 1) NMDG ACSF, 2) SuperACSF, and 3) SuperRecording ACSF. See tables 1-3 for specific notes. All solutions were prepared at room temperature and stored at 4°C. The exact required solutions were made the day prior to use and kept for at most one week or until fully consumed. The preparation follows most of the “protective recovery” methods established in the last decade(13), to both reduce excitotoxicity during decapitation and slicing, as well as supply neuroprotective compounds throughout the entire procedure. The addition of a ketone metabolite is based on literature suggesting the brain preferentially uses ketones (25) during an ischemic state and they may provide a protective effect (26). Carbogenation refers to ultra-fine bubbling of mixed gas (95% O2/5% CO2) into the solutions. Finally, all reagents for solutions were purchased from Sigma.

**Table 1:** Sol1) NMDG ACSF for perfusion, slicing, and the first slice resting period.

| Order | Reagent | mM | MW | g/L |
| --- | --- | --- | --- | --- |
| 1 | NMDG | 92 | 195.2 | **17.96** |
| 2 | KCl | 2.5 | 74.6 | **0.19** |
| 3 | NaH<sub>2</sub>PO<sub>4</sub> | 1.25 | 138.0 | **0.17** |
| 4 | HEPES | 20 | 238.3 | **4.77** |
| 5 | Glucose | 25 | 180.2 | **4.51** |
| 6 | Sodium Ascorbate | 5 | 198.0 | **0.99** |
| 7 | Thiourea | 2 | 76.1 | **0.15** |
| 8 | Myoinositol | 3 | 180.16 | **0.54** |
| 9 | Nicotinamide/B3 | 5 | 122.12 | **0.61** |
| 10 | Taurine | 0.2 | 125.15 | **0.03** |
| 11 | N-Acetyl-L-Cysteine | 12 | 163.19 | **1.96** |
| 12 | Sodium Pyruvate | 3 | 110.0 | **0.33** |
| 13 | β-Hydroxy Butyrate | 1.5 | 126.09 | **0.19** |
| 14 | NaHCO<sub>3</sub> (slowly) | 25 | 84.0 | **2.10** |
| 15 | **pH with HCl first, then NaOH** |  |  |  |
| 16 | CaCl<sub>2</sub>.2H<sub>2</sub>O | 0.5 | 147.0 | **0.07** |
| 17 | MgCl<sub>2</sub> 6H<sub>2</sub>O | 10 | 203.3 | **2.03** |

**Table 2:** Sol2) Super ACSF for the second slice resting period, the recovery & holding period.

| Order |  | mM | MW | g/L |
| --- | --- | --- | --- | --- |
| 1 | NaCl | 92 | 58.4 | **5.38** |
| 2 | KCl | 2.5 | 74.6 | **0.19** |
| 3 | NaH<sub>2</sub>PO<sub>4</sub> | 1.25 | 138.0 | **0.17** |
| 4 | HEPES | 20 | 238.3 | **4.77** |
| 5 | Glucose | 25 | 180.2 | **4.51** |
| 6 | Sodium Ascorbate | 5 | 198.0 | **0.99** |
| 7 | Thiourea | 2 | 76.1 | **0.15** |
| 8 | Myoinositol | 3 | 180.16 | **0.54** |
| 9 | Nicotinamide/B3 | 5 | 122.12 | **0.61** |
| 10 | Taurine | 0.2 | 125.15 | **0.03** |
| 11 | N-Acetyl-L-Cysteine | 12 | 163.19 | **1.96** |
| 12 | Sodium Pyruvate | 3 | 110.0 | **0.33** |
| 13 | β-Hydroxy-Butyrate | 1.5 | 126.09 | **0.19** |
| 14 | NaHCO<sub>3</sub> (slowly) | 25 | 84.0 | **2.10** |
| 15 | **pH with HCl or NaOH as needed** |  |  |  |
| 16 | CaCl<sub>2</sub>.2H<sub>2</sub>O | 2 | 147.0 | **0.29** |
| 17 | MgCl<sub>2</sub> 6H<sub>2</sub>O | 2 | 203.3 | **0.41** |

**Table 3:** Sol3) SuperRecording ACSF for the recording extracellular solution.

| Order of addition |  | mM | MW | g/L |
| --- | --- | --- | --- | --- |
| 1 | NaCl | 125 | 58.4 | **7.31** |
| 2 | KCl | 2.5 | 74.6 | **0.19** |
| 3 | NaH<sub>2</sub>PO<sub>4</sub> | 1.25 | 138.0 | **0.17** |
| 4 | HEPES | 4 | 238.3 | **0.95** |
| 5 | Glucose | 11 | 180.2 | **1.98** |
| 6 | Sodium Ascorbate | 5 | 198.0 | **0.99** |
| 7 | Thiourea | 2 | 76.1 | **0.15** |
| 8 | Myoinositol | 3 | 180.16 | **0.54** |
| 9 | Nicotinamide/B3 | 5 | 122.12 | **0.61** |
| 10 | Taurine | 0.2 | 125.15 | **0.03** |
| 11 | N-Acetyl-L-Cysteine | 6 | 163.19 | **0.98** |
| 12 | Sodium Pyruvate | 3 | 110.0 | **0.33** |
| 13 | β-Hydroxy-Butyrate | 1.5 | 126.09 | **0.19** |
| 14 | NaHCO<sub>3</sub> (slowly) | 24 | 84.0 | **2.02** |
| 15 | **Minor pH adjustment** |  |  |  |
| 16 | CaCl<sub>2</sub>·2H<sub>2</sub>O | 3 | 147.0 | **0.44** |
| 17 | MgCl<sub>2</sub>·6H<sub>2</sub>O | 1 | 203.3 | **0.20** |

The pH should be between 7.3 -7.4, the solution should reach 300-315 mOsm, and ultrapure ddH2O can be used to lower it. Note that on recording days, 5mL of Sol1 should be reserved for a variant solution with 0.58 g of NaCl added; this serves as the intermittent addition of “Na^+^ spike-in” solution during the initial slice resting period. These solutions should be carbogenated at all times. NMDG tends to push the solution to be quite basic, so the major adjustments with several mL of HCl need to be performed prior to minor adjustments with NaOH (if needed).

The pH (7.3 -7.4) and osmolarity (300-315 mOsm) are to be identical to Sol1. Titration is to be done with mostly minor adjustments, as there is no NMDG.

The pH (7.3 -7.4) and osmolarity (300-315 mOsm) are to be identical to Solution 1&2. Titration is to be done with extremely light adjustments, as this solution tends to arrive at nominal pH by default. Note that on recording days, 50mL of Sol3 should be reserved for a variant solution with 0.285 g of KCl added; this serves as the intermittent addition of “75 mM K^+^ spike-in ACSF” solution during the last phase of the recording, to elicit slice-wide activity as a confirmation of health.

### Slice preparation

All animal handling, husbandry, and experimental procedures described were approved by KIST’s Animal Care and Use Committee. Only male transgenic C57BL6 / Thy1-ChR2-YFP mice between 6-10 weeks of age were used, 4 in each group, either corresponding to the 800 µm thick slicing or 400 µm thick slicing. Mice were group housed on *ad libitum* food/water until sacrificed. On the recording days, prior to sacrifice, three variants of Sol1 were obtained: 150 mL of Sol1 was kept in a 400 mL beaker on ice within an insulated vessel, constantly carbogenated for at least 20 min; simultaneously, 250 mL of Sol1 was kept in a 400 mL beaker immersed in a water bath kept at 33°C, constantly carbogenated, and 5 mL “Na^+^ spike-in” solution was made and kept in a 10 mL conical. Next, a 400 mL beaker containing 250 mL of Sol2 was kept in the 33°C water bath, adjacent to the Sol1, constantly carbogenated. Finally, at least 500 mL of Sol3 was kept similarly to Sol2, with 50 mL reserved for mixing the “75mM K^+^ spike-in” ACSF, which was kept in a 50 mL conical. All solutions and the water bath were kept adjacent to a Leica VT1200S vibratome for maximal efficiency. Once all solutions were prepared, 10 mL of the ice-cold, carbogenated Sol1 was loaded into a syringe with a 23g needle, placed within the insulated, ice-filled vessel, and taken with the iced Sol1 to where perfusions were performed; another 70 mL was placed within the ice-covered buffer tray of the vibratome.

Subsequently, a 6-10 week old transgenic mouse was collected and placed within a fume hood for terminal isoflurane anesthesia. Upon cessation of breathing, manual transcardial perfusion with the Sol1 within the syringe was performed, followed by decapitation and placement of the head within the ice-cold Sol1, in total a series of steps taking no more than 2 minutes. Once collected, this Sol1 was re-carbogenated and the head withdrawn after 30 s to ensure adequate cooling. In a series of steps taking less than 3 minutes, the rearmost posterior cranium was cut with surgical scissors, as if to make a coronal section within the posterior cerebellum, and pried off in a rolling motion with surgical forceps, caudal to rostral, gently removing just past the frontal fissure and avoiding puncturing any tissue, after which the brain was then removed by severing attachments to the base of the skull with a surgical spatula, and placed back into the ice-cold Sol1. Within 2 minutes, the center of the Leica specimen disc was dotted with a dime-sized quantity of cyanoacrylate and kept adjacent to a petri dish, within which the excised brain was then placed, whereupon standard single-edged surgical razor blade was used to quickly cut ∼1mm of tissue from the right hemisphere, making a sagittal plane. The brain was transferred using a modified disposable transfer pipette to bring the sagittal (right) plane in contact with the cyanoacrylate; the entire disc was then immersed into the buffer tray, and remaining iced Sol1 was used to submerge the brain as needed, without directly pouring onto the brain. Once positioned with the buffer tray being carbogenated, the entire process would take less than 10 minutes total. The edge of the left hemisphere would be found as the zero-point using the Leica and 410 µm sections would be made until 4100 µm had been cut away, or until the disappearance of the left hippocampus and prior to the appearance of the right. Then, a final 800 µm or 400 µm sagittal slice would be taken depending on the subject. This process would take ∼15 minutes, due to keeping the speed at the lowest setting.

The slice was transferred with a disposable transfer pipette, rather than brushes that pose a risk of damaging slices(20), to a custom loosely mesh-lined holding chamber within the warmed carbogenated 250 mL Sol1, secured with another mesh-lined weight, and according to standard methods, “Na^+^ spike-in” Sol1 was introduced in 5 minute intervals(13). Upon completion of Na^+^ reintroduction in Sol1, the Sol2 beaker was removed from the water bath and the slice was transferred to it in the same way as previously, and held for 1 hour as it recovered and cooled to room temperature, after which it was ready for recording. Note that during the reintroduction and recovery periods, the slices were monitored for bubble formation and removal. For a schematic of the slicing/recording timeline, see Fig. 1f.

### Electrophysiology experiments

Due to supply shortages, both an Intan recording system (RHD2000 USB interface DAQ, Intan Technlogies), and Open Ephys DAQ (Open Ephys Aquisition board, Open Ephys) were used downstream of the C3315 headstages and SPI cables, and an entirely custom data ETL package was written in MATLAB for the Open Ephys DAQ, separate from the commercially available software (Supplemental code database).

Slices ready for recording were held in place using a slice harp on custom mesh webbing suspended within a Petri dish inside a Faraday cage on a vibration table (Supplemental figure 1) to ensure adequate Sol1 flow across the entire slice (21, 27). Room-temperature Sol3 was perfused using gravity and vacuumed off with a pump at a constant, 5-6 mL rate, higher than conventional preparations, to ensure adequate tissue survival(27), with temperature maintained constantly (Warner temperature control system). The 3D probe array was stereotactically lowered to a depth of 600 µm and rested for 10 minutes before recording using a Kopf manual stereotaxis.

Each recording (Fig. 1f) began with 15 minutes of spontaneous activity, followed by two bouts of ChR2 optogenetic stimulation. For the first, a clamped optical fiber from a 473 nm DPSS laser was positioned over the VTA as the landmarks and probe shanks were visible without touching the slice. A trial was initiated with 30 s of quiescence, then pulsed 10 mW/mm^2^ 473 nm light was applied at a 30% duty cycle, .5 Hz in VTA for 100s or at least 30 trials/pulses, and then one minute of quiescence was obtained. Following that, the optical fiber was placed over the mPFC shanks, directly above the slice, and a cortical stimulation was performed, identical to the VTA stimulation. Following that, another 15 minutes of spontaneous activity was recorded, followed by 2mL of 75 mM K^+^ spike-in ACSF addition every 30 seconds (5 second infusion) for 3 minutes. Subsequently, the recordings were terminated and the slice fixed in formalin for imaging at the UNIST Bio-Optics Lab.

### Multiphoton microscopy 3D imaging of the sagittal brain slices

A custom-built multiphoton microscope was used to visualize sagittal brain slices. The multiphoton microscope was equipped with a tunable femtosecond pulse laser (Chameleon Discovery, Coherent) as an excitation source and two photomultiplier tubes (H10770PA-40, Hamamatsu) as detectors. An excitation wavelength of 960 nm was used to excite both YFP and DiI. The detection wavelength range for YFP and Dil was 490 - 560 nm and 570 - 640 nm, respectively using appropriate dichroic and emission filters placed prior to the photomultiplier tubes. A water immersion objective lens (CFI75 LWD 16X W, Nikon) was used to focus the laser and collect the fluorescence signal. For 800 µm thick brain slices, the sample was optically cleared by dipping in an optical clearing solution (C-match, Crayontech) for 12 hours prior to imaging (28) to reduce optical scattering and enable imaging throughout the entire depth. For 400 µm thick slices, imaging was performed on the samples as is. For the whole brain sagittal slice image, the imaging voxel size was 4.6 µm x 4.6 µm x 20 µm (in xyz). A motorized 3-axes translational stage (3DMS, Sutter Instruments) was used to obtain a mosaic of the entire brain slice. The images were stitched using the Stitching plugin in ImageJ. Since different brain regions have different levels of Thy1-YFP expression and neuron density, the excitation laser power was adjusted to obtain enough fluorescence signal in all areas. We also digitally adjusted the contrast for different brain regions to better visualize dim regions. For the high-resolution zoom in images with respect to probe insertion sites, the imaging voxel size was 0.19 µm x 0.19 µm x 2 µm (in xyz). 3D rendering was performed using Fluorender. For a representative image of the 800 µm slice analyzed in Figures 2-4, see Fig. 1g.

**Figure 2.**
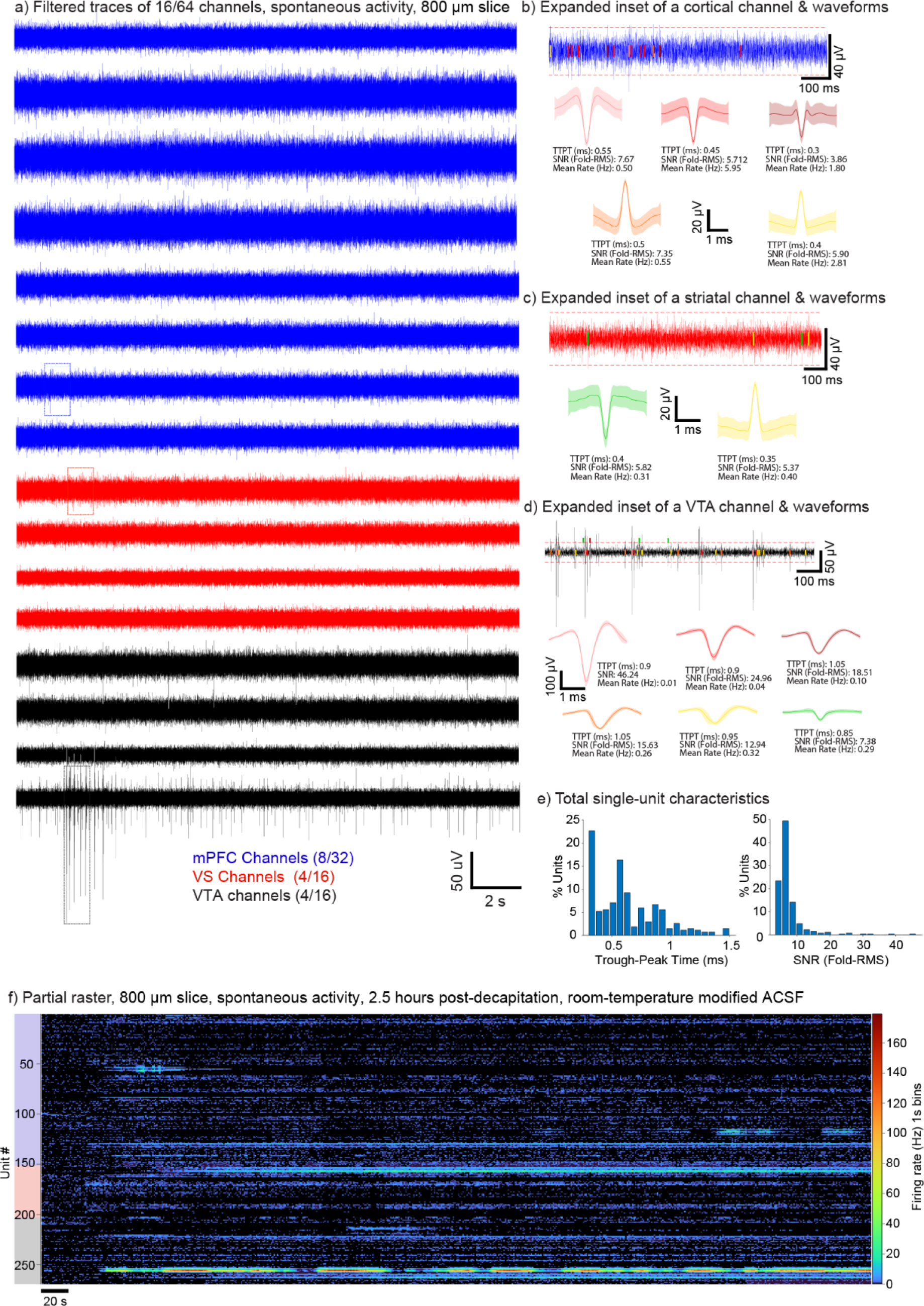
*Spontaneous activity recording from an 800 µm slice.* (a) Filtered channel traces of spontaneous activity with region and color indicated for consistency; insets indicate timeseries for (b), (c), and (d) for the appropriate regions. Single unit waveforms with shaded standard deviation are color-coded to associate within their panel inset; red dashes indicate threshold for detection. TTPT = trough-peak time, SNR = signal to noise ratio. Indicated for each waveform are a subset of the descriptive statistics collected for each single unit. (e) Trough-peak time (left) and SNR (right) distributions as histograms for one slice. Note the bimodal distribution for TTPT, consistent with *in vivo* recordings. (f) Partial activity raster for one slice with unit location indicated as color on left edge.

### Electrophysiology data analysis

Due to the usage of custom silicon polytrode arrays and *ex vivo* recording of signals, a highly-optimized custom spike-sorting algorithm was developed to examine the low-noise, low signal neural features, as other available algorithms failed to detect signals (29), failed to present an efficient pipeline and had far too many dependencies (2), or required certain assumptions about the electrode hardware and recording software. Thus, we present a semi-automated solution and platform for *ex vivo* analysis that grants the end-user full control of the data at every step, with clear instructions (supplemental code).

Briefly, data were sampled at 20 kHz under a 4 pole Butterworth filter, with a passband of 300-6000 Hz and any channels with excessive 60 Hz noise in the raw trace were omitted from analysis. In MATLAB, raw voltage data were subject to a cascading, 5-harmonic notch filter based around 60 Hz, subjected to the same filter as during the recording, and S-G smoothed with amplitudes corrected. Threshold was set to be 5x RMS of each channel’s trace (30) and upon visual inspection of the trace, a per-subject correction was applied to capture obvious deflections, lowering threshold by up to 15%.

Putative times were binned into a logical matrix with 1 ms windows and overlapping proto-spiketime bins were pruned if appearing on all channels across the array, thus removing ISI violations, laser artefacts, and table motion/other motion artefacts. Proto-waveforms were associated with those times, and the appearance of an overlapping timestamp within a two-shank neighborhood, with a waveform of the same sign, within 0.2 ms, would cause for adjudication between presenting channels based on amplitude. We reason that due to the overall sparseness of neural signals(9, 31) and absence of a significant amount of LFP noise *ex vivo*(*32*), the spacing between recording sites, and how action potentials (APs) decay through tissue(33), signals can be reliably parsed and assigned to adjacent channels consistently. Final spike times were associated with 3.2 ms filtered waveforms, with subsequent PCA and Z-Scoring performed on all waveforms. Waveforms underwent K-Means over-clustering with an elbow-method for automatic K determination, with an assisted merging step, across all channels (Supplemental Fig. 2). Finally, single units underwent semi-automatic unit quality assessment, a battery of descriptive statistics (Fig. 2b-d, explanations below each mean unit trace), and in the case of optogenetics experiments, a paired T-test with rates compared between laser stimulation periods (during the laser and 1.5 laser-length periods afterwards) and the pre-laser intertrial periods (two laser lengths prior to onset). LFP data were taken from the raw data, per channel, through downsampling to 1 kHz, after the notch filtering step. Due to the intrinsic lack of excess noise in this recording platform, no background subtraction was necessary.

### Statistical Testing

All statistical analysis was performed with standard MATLAB functions or GraphPad Prism software and tests are indicated in-line and within legends as indicated.

## Results

### Spontaneous activity recording from an 800 µm slice

The following represents a summary of data collected from one representative C57BL6 / Thy1-ChR2-YFP animal. To our knowledge, since no group thus far has demonstrated the electrophysiological properties of brain slices more than twice the thickness of conventional slices, and since the general consensus of the ischemia and hypoxia literature suggests that thick brain slices suffer necrosis internally, direct observation of the deep interior of the slice’s electrophysiological properties during acute experimentation can both yield more insight about the timing of ischemia, as well as provide evidence for the viability of this platform as an alternative method to work on intact circuits(18–21).

Qualitatively and quantitatively, a wide variety of single-unit responses can be obtained across the mPFC (Fig. 2a, blue traces & inset for Fig. 2b), VS (Fig. 2a, red traces & inset for Fig. 2c), and VTA (Fig. 2a, black traces & inset for Fig. 2d). Additionally, our single slice distribution of Trough-to-Peak Times (Fig. 2e, left) and amplitude expressed as multiples of SNR (Fig. 2e, right) are consistent with canonical multiregion *in vivo* recordings(1, 34) suggesting the inherent viability of the technique, with the added benefit of being able to detect units at a signal threshold of 5x the background RMS band of 5.56±2.08 µV, orders of magnitude lower than typical *in vivo* recordings, similar to organoids; strikingly, this is near the 2.4 μV noise floor of the Intan chips themselves (Intan Technologies materials). This also allows us to achieve unit yields multiple times greater than current high-density neuroprobe methods with redundant channels(34, 35). Finally, conventional neurophysiological knowledge suggests that positive-deflecting spikes are only a small percentage of single-unit responses, but in our hands, nearly 40% of the units we record are based on positive spikes, as in Figures 2b & 2c.

Additionally, we do not see a correlation between depth of channel, firing rate, and duration of the recording in this representative animal (Supplemental Fig. 3) and these correlations were near zero, suggesting that further tests in other animals, given the number of units, duration bins, and depths, would also be close to zero. This demonstrates that nearly 2.5 hours after initial decapitation, our *ex vivo* preparation remains robust and has *in vivo-like* activity dynamics (Fig. 2f), while displaying scant direct physiological correlates of ischemia or hypoxia. Anecdotally, in this slice, some of the highest firing rates were observed close to the middle of the shanks, in the core of the tissue (Supplemental Fig. 3). Finally, the overall long-tailed distribution of firing rates in this slice is highly consistent with evidence supporting the “dark neuron” problem of a general lack of robust responses *in vivo* (Supplemental Fig. 4) (36), which is typically not discussed, rather, these units tend to be ignored to support experimenter-led claims. This is further evidence in favor of the more physiological nature of our preparation and its suitability for further experimentation as a platform.

### Characterization of optogenetic stimulation in VTA from an 800 µm slice

To demonstrate the functionality of the MFB in an acute, sagittal, tract-preserving thick slice several hours after decapitation upon completion of spontaneous activity recording, ChR2- based firing was evoked in the VTA in order to elicit firing throughout the mesocorticolimbic circuit *vis a vis* disinhibition(37), reinforcing facilitation(38), and putative glutamate co-release(39) (Fig. 3a). Binned single-unit firing qualitatively indicated robust firing response when the laser was pulsed, particularly within VTA units (Fig. 3b, units 100-165, interleaved with VS units). A subset of simultaneous VTA (Fig. 3c, top two black traces), VS (Fig. 3c, middle two red traces), and mPFC (Fig. 3c, blue trace) channels are indicated with stimulation times.

**Figure 3.**
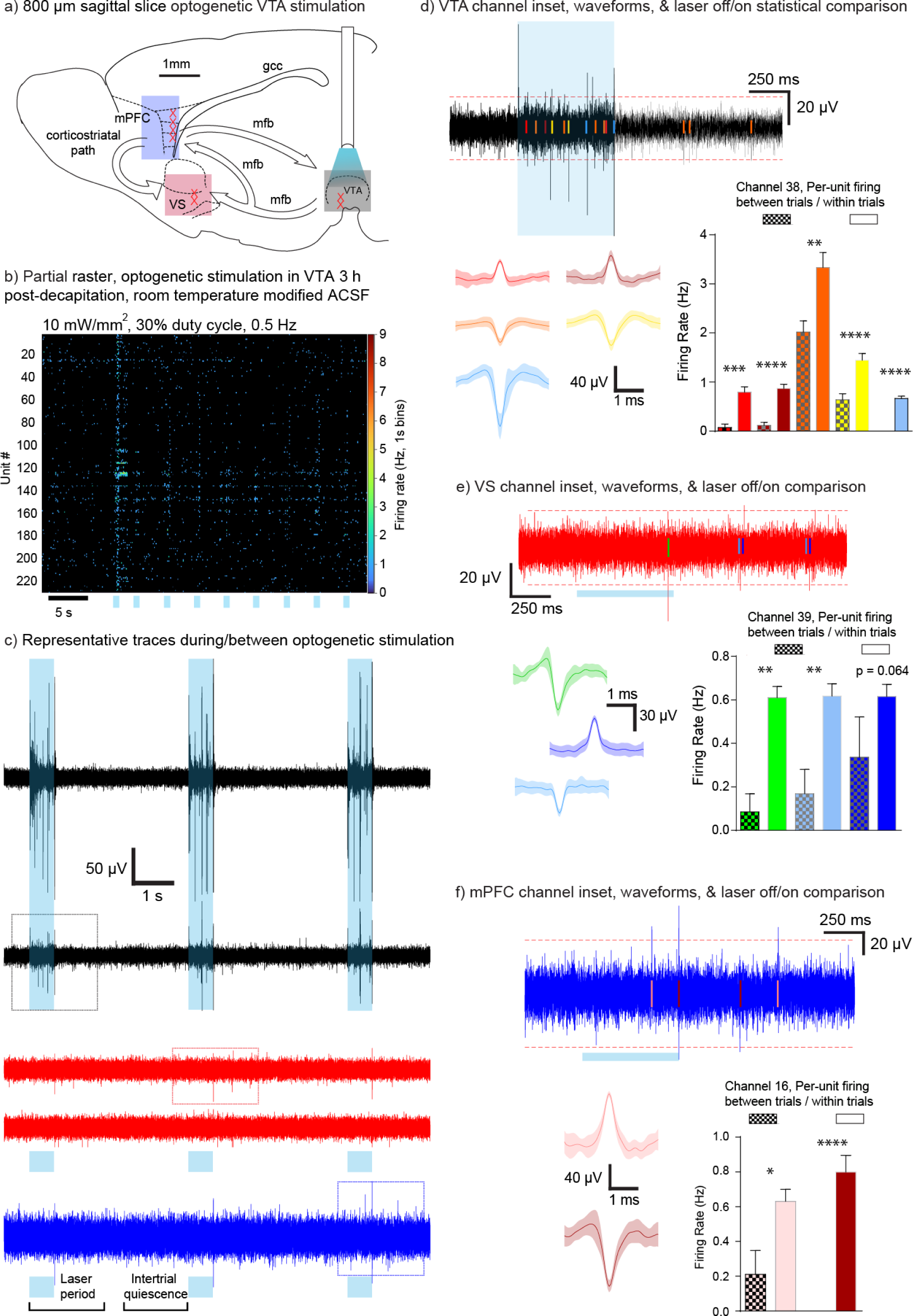
*Characterization of optogenetic stimulation in VTA from an 800 µm slice.* (a) Stimulation schematic of the slice, where VTA stimulation feeds activity into the VS and mPFC. (b) Partial firing raster with laser pulses indicated. (c) Simultaneous filtered voltage traces of a subset of channels per region. Note that the laser stimulation period includes 1.5 laser pulse widths worth of time after the laser ends to capture delayed activity, while the intertrial quiescent period is two pulse widths long, prior to next laser onset. (d)-(f) Insets from (c), per region, color- coded single unit waveforms with shaded standard standard deviation. Each single unit’s firing rate underwent a paired T-test between laser trial and intertrial periods, on the bottom right, with significance levels below p = .05 indicated as asterisks; otherwise exact p indicated.

Threshold-passing events are time-locked to the stimulation period, across the channels, at the same time, indicating a functional circuit may be present.

Examining each trace in detail yields a variety of unit responses within the stimulation region of the VTA (Fig. 3d, black inset from Fig. 3c). In the represented channel (with thresholds indicated as red-dashes), at the indicated timeseries within the inset, all the color-coded single-units have a highly statistically significant elevation of firing rate over the intertrial period with all two-tailed, paired T-tests surpassing p < .01 (Fig. 3d, bottom right). This indicates that at a subsecond scale and across dozens of trials, throughout the entire depth of the recording electrodes, neural tissue remained robust enough to be responsive to the stimulation.

Examining the projection targets of the VTA, the VS (Fig. 3e, red inset from Fig. 3c) and mPFC (Fig. 3f, blue inset from Fig. 3c) shows fewer per-channel single-units, but each channel does have a subset of responsive units according to the within trial vs. between trial criteria. This demonstrates that some degree of optogenetically-evoked firing can confidently be observed in thick sagittal slices of murine tissue, potentially based on the integrity of the MFB.

### Characterization of optogenetic stimulation in mPFC from an 800 µm slice

If the MFB is intact in our preparation, it stands to reason that identical stimulation of the cortical inputs (Fig. 4a) should likewise elicit some degree of glutamatergic-based excitation throughout the other hubs within the mesocorticolimbic circuit. As in the VTA previously, binned single-unit firing qualitatively indicated robust firing response when the laser was pulsed, particularly within mPFC units (Fig. 4b, units 1-69 & 157-200). A subset of simultaneous mPFC (Fig. 4c, top two blue traces), VS (Fig. 4c, middle two red traces), and VTA (Fig. 4c, black trace) channels are indicated with stimulation times. Threshold-passing events are time-locked to the stimulation period, across the channels, at the same time, indicating a functional circuit may be present.

**Figure 4.**
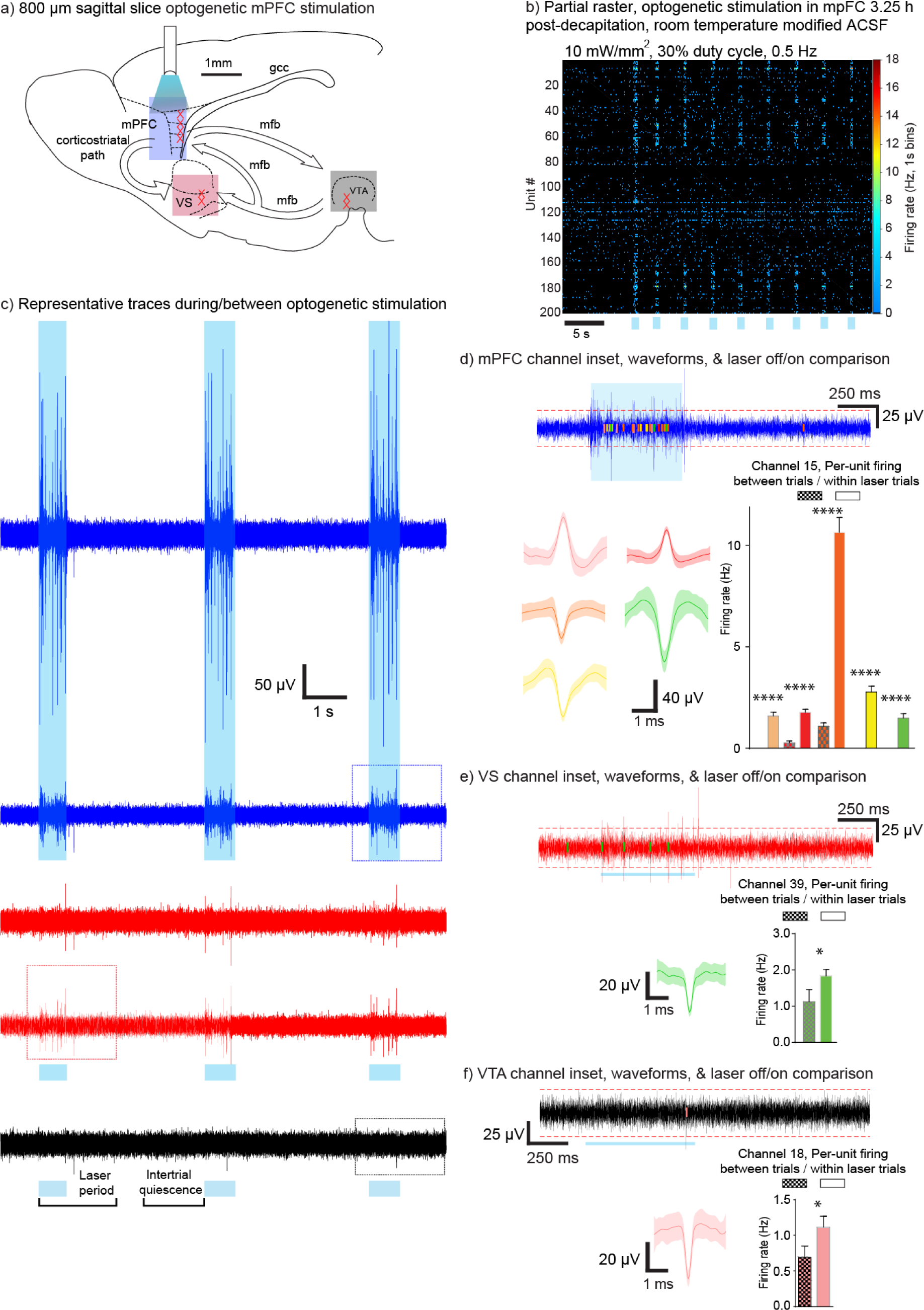
*Characterization of optogenetic stimulation in mPFC from an 800 µm slice.* (a) Stimulation schematic of the slice, where mPFC stimulation feeds activity to VS and Vta. (b) Partial firing raster with laser pulses indicated. (c) Simultaneous filtered voltage traces of a subset of channels per region, as in Figure 3. (d)-(f) Insets from (c), per region, color-coded single unit waveforms with shaded standard standard deviation. Each single unit’s firing rate underwent a paired T-test between laser trial and intertrial periods, on the bottom right, with significance levels below p = .05 indicated as asterisks.

Further inspection of the mPFC again shows a variety of unit responses within the stimulation region (Fig. 4d, blue inset from Fig. 4c). Again, in the represented mPFC channel, all the color-coded single-units have a very highly statistically significant elevation of firing rate over the intertrial period with all two-tailed, paired T-tests surpassing p < .0001 (Fig. 4d, bottom right). As previously, the other mesocorticolimbic hubs we recorded also have statistically significant evoked activity within the laser trial periods vs. intertrial quiescence (Fig. 4e, red inset from Fig. 4c & Fig. 4f, black inset from Fig. 4c), and similarly, the number of units/channel is lower at the projection targets. Having demonstrated the viability of both spontaneous and long-range evoked activity in an acute thick brain slice, as well as our analysis pipeline for efficient reporting of unit statistics, we next sought to ascertain if the thicker brain slice would have a more robust functional coupling between regions. This would succinctly demonstrate the viability and advantages of using thick brain slices for circuit mapping and pharmacological testing *ex vivo*, and to that end, we performed the previous experiments across four 800 µm and four 400 µm C57BL6 / Thy1-ChR2-YFP sagittal slices.

### Thick slices preserve asymmetric communication across the mesocorticolimbic circuit

Given that we can record hundreds of single units per animal, the statistical power we generate when optogenetically stimulating needs to be leveraged with group and population-based metrics(40). The simplest and most robust mathematical approach is to consider the count of statistically laser-responsive neurons per region (examples seen in Figures 3 & 4), rather than dealing with correlation coefficients or other fine-scale metrics of neural functional connectivity that are spurious and taper off(41). As we are comparing different numbers of units overall in each slice condition, the count is turned into a proportion, and we’ve designated this metric as ε.

When considering ε in VTA units responding to VTA stimulations, across animals and slice conditions, there was no statistically discernible difference between the 800 µm and 400µm slices (Fig. 5a, left, one-tailed Mann-Whitney U test, p = .329), and given that there was identical optogenetic drive in all conditions, this would be consistent with equal magnitude of local response. However, within the VS, we observed an enhanced ε in 800 µm slices (Fig. 5a, middle, one-tailed Mann-Whitney U test, p = .043) as compared to 400 µm slices. Similarly, this enhancement in thicker slices was more apparent in the mPFC (Fig. 5a, right, one-tailed Mann-Whitney U test, p = .014). This is consistent with the hypothesis that a thicker brain slice would be more functionally connected due to retaining more of the MFB.

**Figure 5.**
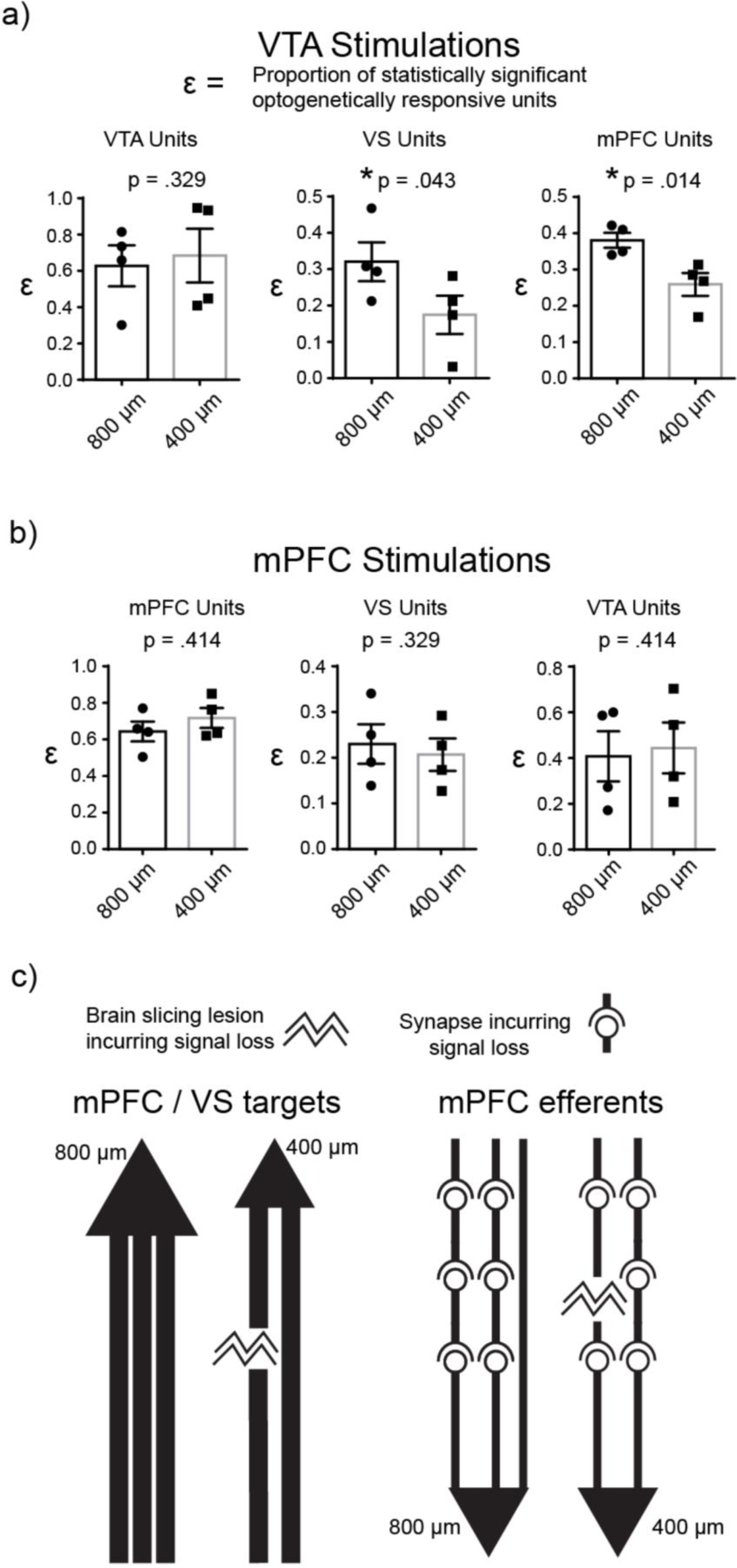
*Thick slices preserve asymmetric communication across the mesocorticolimbic circuit* ε = proportion of statistically optogenetically-responsive neurons per region. (a) Within VTA stimulations, comparing ε between slice types among VTA units revealed no statistically discernible difference (left, one-tailed Mann-Whitney U test, p = .329) likely due to identical optogenetic drive, but a difference was found in the VS (middle, one-tailed Mann-Whitney U test, p = .043) and mPFC (right, one-tailed Mann-Whitney U test, p = .014), with the thicker slices in general having a greater proportion of units firing within the laser trial periods. (b) Within mPFC stimulations, no brain region had a statistically discernible difference in ε slices (mPFC, left, one-tailed Mann-Whitney U test, p = .414; VS, middle, one-tailed Mann-Whitney U test, p = .329; VTA, right, one-tailed Mann-Whitney U test, p = .414). (c) The preceding results are consistent with the idea of an asymmetric bidirectionality among the hubs in the mesocorticolimbic circuit carried by the MFB, illustrated in this schematic. Monosynaptic VTA efferents take a noticeable loss in functional connectivity after severing a greater amount in the 400 µm condition (left), but multisynaptic mPFC efferents, already having loss in synapses when reaching targets, have less detectable difference in functional connectivity after increased severing during production of 400 µm slices (right).

As before in the VTA, mPFC units responding to local cortical stimulation had identical optogenetic drive across slice conditions and had no statistically discernible difference in ε between the 800 µm and 400 µm slices (Fig. 5b, left, one-tailed Mann-Whitney U test, p = .414). Surprisingly, we did not see an enhancement in ε for the thicker slices neither in the VS (Fig. 5b, middle, one-tailed Mann-Whitney U test, p = .329) nor in the VTA (Fig. 5b, right, one-tailed Mann-Whitney U test, p = .414).

The asymmetric bidirectionality of this circuit can be readily assessed *ex vivo* in thick slices without complicated surgical procedures and implants, and we hypothesize that according to established anatomy, our result fits into canonical understanding of the MFB. Specifically, the disinhibitory/reinforcing(37, 38) dopaminergic and occasionally glutaminergic projections from the VTA(39) are primarily monosynaptic onto the VS and mPFC (42). Thus, they are severed through conventional thin slices or coronal sections and would have lower functional connectivity, as in ε (Fig. 5c, left). However, mPFC efferents are mired in denser local circuitry, and it has been known for decades in addiction research that mPFC efferents arriving in VTA are trans-synaptic(16). Thus in our schematic, even if one severs tracts by cutting a thinner section, the more intact preparation still has many synapses where local inhibition or basic signal propagation principles can prevent APs from crossing synapses, making the end results look similar for mPFC stimulations (Fig. 5c, right).

## Discussion

Combining a suite of custom technical polytrode features, essential neurophysiological principles, disparate but complementary approaches in electrophysiology, and a highly tailored data analysis approach, we have developed a platform and protocol for neural recordings that combines the best features of silicon probe electrophysiology *in vivo* with those of *ex vivo* preparations. At first glance, our result being highly consistent with established literature is not surprising, but underpinning it is a protocol that has never been attempted due to requiring different probes, full understanding of slice physiology, and the data fluency of high-density recordings.

We first demonstrated that our novel technology and recording pipeline could generate *in vivo-*like activity patterns across three hubs in a highly-druggable circuit using thick brain slices that would be difficult, if not impossible to record *in vivo*. This activity was robust hours after brain extraction, a novel result indicating the current inadequacies in hypoxia and ischemia literature (19, 20, 22, 23). Our methodology revealed consistencies with other *ex vivo* recording methods, such as very low noise (32, 43), that, taken with greater channel spacing, allows for a greater tissue coverage and many times greater yield per channel of our method over current silicon probes, all while running on easily available hardware and a software package with no dependencies. By showing that thicker slices have functional circuits with greater connectivity due to retaining tracts, expanding upon experiments using more functional slices(39), we also verified that these connections are not symmetrical, and this platform represents a viable potential methodology to approach pharmacological mechanism of action (MoA) studies in conjunction with patch clamp by providing extra physiological and translational relevance with no cost to the experimenter.

One drawback of our ε metric is that it may be argued to be too simple; yet that is its chief advantage, given that the question we ask is simple. Other metrics such as correlations require fine-tuning of temporal windows and require higher firing rates, or others such as mutual information, Granger causality (44), and graph theory metrics are computationally more demanding and difficult to explain to a wide audience. Another issue potentially is that our number of mice in each group is only four, requiring the usage of non-parametric statistics, yet our reasoning is twofold. We perform research in best accordance with the “3 Rs” principle and maximize animal welfare where possible. Additionally, this implicitly helps strengthen the credibility of the result - if it were not robust with such a low number of animals and large number of units behind each animal, boosting the number would be akin to p-hacking and not a truly scientifically robust result.

It is widely assumed that the utility of neurophysiology in understanding neural encoding or pharmacological MoA lies at the two extremes of experimental preparations: high-density *in vivo* and whole-cell patch clamp, respectively, with the drawbacks of each minimized in their discussion. It should be noted that adherence to such dichotomous preparations has yielded methodology for understanding experimentation orthogonal to practical questions, since many assumptions underlying each are rarely critically discussed despite being *prima facie* contrived to the point of esotericism. The current push to use alternative models such as organoids or pure *in silica* models for drug testing or basic science is yet even a further step of artificiality which engenders not only epistemological, but also ontological and pragmatic risk.

For example, a robust, reproducible finding is that *in vivo* extracellular recordings and calcium imaging datasets typically yield active signaling unit equivalents at a rate of at least one order of magnitude lower than would be expected via anatomical estimates(31, 33). Not only that, but *in vivo* RMS noise tends to be upwards of 40 μV, drastically limiting the number of units discoverable, due to the LFP being a summation of all electrical signals at the location one has implanted an electrode. The resultant neurons of focus, “non-dark” and “type 2 dark” neurons end up skewing all analyses(36) based on population averages, if even the murine or human brain “cares” about arbitrary average firing rates within experimenter-defined windows(45). Ultimately this “sparse” but skewed activity representation *in vivo* can generate essentially statistical artifacts of “encoding” such as “grandmother”-like cells (46), units explicitly tuned to concepts like numbers (47), and brain-wide reward-responsive neurons. This is a certainty known to be a byproduct of enhanced statistical power due to the vastly increased yields of newer high-density recording methods coupled to older frequentist statistics. Even if neural activity patterns were not biased by experimenter-chosen analysis pipelines or ignoring largely quiescent “dark” neurons, the issue of murine behavior and the effects thereon by implants & surgery, housing & handling (48), task structure, diet, and inbreeding & enhanced telomere length *vis a vis* the implications of antagonistic pleiotropy (49), is frequently not discussed and assumed to have little to no effect on neural activity patterns or potentially compensatory complex responses to experimental perturbations. *In vivo* behavior, despite all these confounds or lurking variables, is still a gold standard for efficacy of pharmacological testing or algorithmically-led neural decoding, which could also represent a principal component of the failure to bring, for example, Alzheimer’s drugs to market among nearly all pharmaceutical companies for decades despite billions of dollars of funding (50), and a lack of reproducibility in basic science (51).

Our extremely low-noise recording platform with large electrode coverage both observes “dark neurons” and signals distal to recording sites, extending the practicality of lower-density polytrode arrays, while defining “ground truth” in recording data, due to not arbitrarily throwing out units that do not satisfy investigator activity criteria(36). Such enhancement in the utility of recording technology can go hand-in-hand with AI-based machine-learning methods to data-mine the patterns in the slice, and even work synergistically with behavior and data collected prior to sacrifice. Ideally, extremely miniscule changes in physiology are now more observable due to this technique, allowing for small modifications in drug development and assays to target particular circuits.

On the other hand, glass pipette-based physiology also has a bias towards monitoring visibly healthy neurons while effectively reducing channel count to one or at most a handful of neurons entirely dependent on experimenter skill(14). Such granularity is important strictly for ascertaining channel functionality but of exceedingly limited use in deducing mesoscale connectivity; thus in particular, this technique has remained the workhorse of pharmacological testing in slice or culture. Yet, no slice or tissue culture or experiments can yield a physiological multi-region circuit, thus drastically limiting conclusions about connectivity and pharmacological effects, especially considering that it is not brain slices, but human patients, that take medications. Our method allows for a correction in this flawed preparation and, although we have not yet attempted day-long recordings as is conventional for slice protocols, we are confident that the data and methods reported here will allow for those tests to be forthcoming. Additionally, further experimentation will help to determine the upper limits on tissue thickness, as it will be unlikely that whole-brain samples will be as robust (52). Even so, evidence from months-old organoids with a diameter of 1mm suggests that a necrotic, hypoxic core is not guaranteed even in that condition (53, 54). Thus, measuring extremely minute and distant electrophysiological changes in a 1mm or more thick slice may offer a more realistic alternative to the artificial networks and seizure-like “burst or bust” activity apparent in neural organoids(32).

There are a few biophysical curiosities of note in our experiments: the amplitude of our units tended to be lower than those observed *in vivo*, the optogenetic activity propagation tended to be slow and generally weak which may also lead to questions about the interpretation of Figures 3-5, and we have seen intracellular EPSPs with ostensibly extracellular electrodes (Supplemental figure 5). Regarding the amplitude of our spikes, this is a commonality with both organoids(32) and 2D MEA culture experiments(43), and thus may be unique to non-*in vivo* setups. Knowing that only a fraction of neurons present within the physical AP-detection window around a recording channel will actually fire during recording epochs(9, 31), and knowing that each channel’s ability to detect decreases as a function of distance from the putative unit(31), it becomes a mathematical certainty that the AP amplitudes detected will follow a distribution, and most will fall along the left side of it (Fig. 2e, right)(1). Additionally, it is known that recording at lower temperatures as we have done in order to keep tissue healthier for longer(55) may also result in lower amplitude spikes(56–58).

As for the amount of optogenetic activity we recorded, there is a known feature of using long stimulation periods in Thy1-ChR2 mice, in which depolarization block occurs alongside recruitment of inhibitory currents up to hundreds of milliseconds after initiation of the pulse(59). This can explain in general both low firing rates and variability in responses, as all neurons local to the stimulation, including abundant inhibitory neurons that we can’t record, are recruited.

Optogenetic stimulation of the VTA is also known to take effect in VS even several hundreds of milliseconds after pulses initiate under anesthesia(60), establishing that even in monosynaptic circuits, delays on that order are physiological, hence why we considered trial lengths to be more than one second long. We acquiesce that it may be possible that lowering the stimulation time would counterintuitively increase the fidelity of our evoked signals, and will consider that an avenue for future experimentation.

Finally, it is an uncommon but not impossible occurrence for us to record EPSPs as if we had obtained a gigaseal, as this happened in one out of eight of our slices, but represents an opportunity to understand how to better our techniques to optimize for this high-granularity information. Others have seen this similar intracellular-like data with tetrodes(61) and cultured MEA methods (62, 63), and it stands to reason that this is an outcome far more common than is conventionally thought, but only visible with the right analysis pipeline.

In summary, we demonstrate a method to record *ex vivo* deep-brain functional connectivity which may provide experimenters a platform for acute circuit manipulations beyond what is available in certain in vivo settings, bridging the gap between *ex vivo* and *in vivo* research. This represents a more realistic and physiological circuit-based target for *ex vivo* drug testing due to the ability to finely alter and record many more previously undetectable signals from the slice, and has many open paths of further experimentation. These include, but are not limited to, assays of transplanted and excised tissue, real-time multifiber circuit manipulation coupled to pharmacological testing, and in-depth neurophysics methods(35) and machine learning to characterize the slices in a manner heretofore unseen. There are yet unknown aspects to the progression of many neurodegenerative diseases which may become druggable targets, and this represents a strong method and platform to grasp the circuit and population correlates of established pharmacodynamics or the potentially silent phenotypes therein.

## Acknowledgements

This work was supported by the National Research Foundation (NRF) Brain Pool Program through the Korean Federation of Science and Technology Societies (KOFST) funded by the Ministry of Science ICT and Future Planning (Grant 2021H1D3A2A02082997). Additionally, it was supported by the Brain Convergence Research Program of the National Research Foundation (NRF) funded by the Korean government (MSIT) (NRF-2019M3E5D2A01063814) and Research program for understanding and regulation of brain function of the National Research Foundation (NRF) funded by the Korean government (MSIT) (NRF-2022M3E5E8081196). The authors would like to thank all the generous, helpful members of the KIST community including the Nakwon Choi lab, Bradley Baker Lab, and Sébastien Royer Lab for their advice and suggestions.

## Author contributions

WCS designed and performed research, contributed new protocols/analytic tools and designs, analyzed data, made figures, and wrote the paper. ZN contributed new materials, probe designs, and made figures. HS & UC contributed materials and probes. SHL & JHP performed tissue imaging and made figures. YDC, JWW, SKK, SJO, & MHN provided materials, reagents, and equipment. TSK & IJC designed research, analyzed data, and edited the manuscript.

## References

1. Shobe JL, Claar LD, Parhami S, Bakhurin KI, & Masmanidis SC (2015) Brain activity mapping at multiple scales with silicon microprobes containing 1,024 electrodes. Journal of neurophysiology 114(3):2043–2052.

2. Steinmetz NA, Koch C, Harris KD, & Carandini M (2018) Challenges and opportunities for large-scale electrophysiology with Neuropixels probes. Current opinion in neurobiology 50:92–100.

3. Saxena T, et al. (2013) The impact of chronic blood-brain barrier breach on intracortical electrode function. Biomaterials 34(20):4703–4713.

4. Todorov MI, et al. (2020) Machine learning analysis of whole mouse brain vasculature. Nature methods 17(4):442–449.

5. Papaioannou S & Medini P (2022) Advantages, Pitfalls, and Developments of All Optical Interrogation Strategies of Microcircuits in vivo. Frontiers in neuroscience 16:859803.

6. Okun M, Lak A, Carandini M, & Harris KD (2016) Long Term Recordings with Immobile Silicon Probes in the Mouse Cortex. PloS one 11(3):e0151180.

7. Chae U, et al. (2024) KDS2010, a reversible MAO-B inhibitor, extends the lifetime of neural probes by preventing glial scar formation. Glia.

8. Gaire J, et al. (2018) The role of inflammation on the functionality of intracortical microelectrodes. Journal of neural engineering 15(6):066027.

9. Shoham S, O’Connor DH, & Segev R (2006) How silent is the brain: is there a “dark matter” problem in neuroscience? Journal of comparative physiology. A, Neuroethology, sensory, neural, and behavioral physiology 192(8):777–784.

10. Scanziani M & Hausser M (2009) Electrophysiology in the age of light. Nature 461(7266):930–939.

11. Piatkevich KD, et al. (2019) Population imaging of neural activity in awake behaving mice. Nature 574(7778):413–417.

12. Schmidt-Supprian M & Rajewsky K (2007) Vagaries of conditional gene targeting. Nature immunology 8(7):665–668.

13. Ting JT, Daigle TL, Chen Q, & Feng G (2014) Acute brain slice methods for adult and aging animals: application of targeted patch clamp analysis and optogenetics. Methods in molecular biology 1183:221–242.

14. Obermayer J, et al. (2018) Lateral inhibition by Martinotti interneurons is facilitated by cholinergic inputs in human and mouse neocortex. Nature communications 9(1):4101.

15. Rock KL & Kono H (2008) The inflammatory response to cell death. Annual review of pathology 3:99–126.

16. Wise RA (2005) Forebrain substrates of reward and motivation. The Journal of comparative neurology 493(1):115–121.

17. Arenkiel BR, et al. (2007) In vivo light-induced activation of neural circuitry in transgenic mice expressing channelrhodopsin-2. Neuron 54(2):205–218.

18. Wang T & Kass IS (1997) Preparation of brain slices. Methods in molecular biology 72:1–14.

19. Newman GC, Hospod FE, & Wu P (1988) Thick brain slices model the ischemic penumbra. Journal of cerebral blood flow and metabolism : official journal of the International Society of Cerebral Blood Flow and Metabolism 8(4):586–597.

20. Papouin T & Haydon PG (2018) Obtaining Acute Brain Slices. Bio-protocol 8(2).

21. Killian NJ, Vernekar VN, Potter SM, & Vukasinovic J (2016) A Device for Long-Term Perfusion, Imaging, and Electrical Interfacing of Brain Tissue In vitro. Frontiers in neuroscience 10:135.

22. Voss LJ (2020) Relationship between artificial cerebrospinal fluid oxygenation, slice depth and tissue performance in submerged brain slice experiments. Neuroscience letters 736:135275.

23. Hrabetova S & Nicholson C (2000) Dextran decreases extracellular tortuosity in thick-slice ischemia model. Journal of cerebral blood flow and metabolism : official journal of the International Society of Cerebral Blood Flow and Metabolism 20(9):1306–1310.

24. Shin H, et al. (2019) Multifunctional multi-shank neural probe for investigating and modulating long-range neural circuits in vivo. Nature communications 10(1):3777.

25. White H & Venkatesh B (2011) Clinical review: ketones and brain injury. Critical care 15(2):219.

26. Makievskaya CI, et al. (2023) Ketogenic Diet and Ketone Bodies against Ischemic Injury: Targets, Mechanisms, and Therapeutic Potential. International journal of molecular sciences 24(3).

27. Hajos N, et al. (2009) Maintaining network activity in submerged hippocampal slices: importance of oxygen supply. The European journal of neuroscience 29(2):319–327.

28. Kim K, et al. (2022) Optimized single-step optical clearing solution for 3D volume imaging of biological structures. Communications biology 5(1):431.

29. Swindale NV, Mitelut C, Murphy TH, & Spacek MA (2017) A Visual Guide to Sorting Electrophysiological Recordings Using ’SpikeSorter’. Journal of visualized experiments : JoVE (120).

30. Chaure FJ, Rey HG, & Quian Quiroga R (2018) A novel and fully automatic spike-sorting implementation with variable number of features. Journal of neurophysiology 120(4):1859–1871.

31. Henze DA, et al. (2000) Intracellular features predicted by extracellular recordings in the hippocampus in vivo. Journal of neurophysiology 84(1):390–400.

32. McDonald M, et al. (2023) A mesh microelectrode array for non-invasive electrophysiology within neural organoids. Biosensors & bioelectronics 228:115223.

33. Quiroga RQ (2012) Spike sorting. Current biology : CB 22(2):R45–46.

34. Smith WC, et al. (2016) Frontostriatal Circuit Dynamics Correlate with Cocaine Cue-Evoked Behavioral Arousal during Early Abstinence. eNeuro 3(3).

35. Smith WC (2022) In vivo Quantification of Neural Criticality and Complexity in Mouse Cortex and Striatum in a Model of Cocaine Abstinence. *bioRxiv*:2022.2008.2002.501652.

36. Wohrer A, Humphries MD, & Machens CK (2013) Population-wide distributions of neural activity during perceptual decision-making. Progress in neurobiology 103:156–193.

37. Ambrosi P & Lerner TN (2022) Striatonigrostriatal circuit architecture for disinhibition of dopamine signaling. Cell reports 40(7):111228.

38. Fisher SD, et al. (2017) Reinforcement determines the timing dependence of corticostriatal synaptic plasticity in vivo. Nature communications 8(1):334.

39. Chuhma N, Choi WY, Mingote S, & Rayport S (2009) Dopamine neuron glutamate cotransmission: frequency-dependent modulation in the mesoventromedial projection. Neuroscience 164(3):1068–1083.

40. Kohn A, Coen-Cagli R, Kanitscheider I, & Pouget A (2016) Correlations and Neuronal Population Information. Annual review of neuroscience 39:237–256.

41. Cohen MR & Kohn A (2011) Measuring and interpreting neuronal correlations. Nature neuroscience 14(7):811–819.

42. Oh SW, et al. (2014) A mesoscale connectome of the mouse brain. Nature 508(7495):207–214.

43. Radivojevic M & Rostedt Punga A (2023) Functional imaging of conduction dynamics in cortical and spinal axons. eLife 12.

44. Trongnetrpunya A, et al. (2015) Assessing Granger Causality in Electrophysiological Data: Removing the Adverse Effects of Common Signals via Bipolar Derivations. Frontiers in systems neuroscience 9:189.

45. Tlaie A, Shapcott KA, Tiesinga P, Schölvinck ML, & Havenith MN (2021) Does the brain care about averages? A simple test. *bioRxiv*:2021.2011.2028.469673.

46. Barwich AS (2019) The Value of Failure in Science: The Story of Grandmother Cells in Neuroscience. Frontiers in neuroscience 13:1121.

47. Kutter EF, et al. (2023) Distinct neuronal representation of small and large numbers in the human medial temporal lobe. Nature human behaviour 7(11):1998–2007.

48. Sensini F, et al. (2020) The impact of handling technique and handling frequency on laboratory mouse welfare is sex-specific. Scientific reports 10(1):17281.

49. Weinstein BS & Ciszek D (2002) The reserve-capacity hypothesis: evolutionary origins and modern implications of the trade-off between tumor-suppression and tissue-repair. Experimental gerontology 37(5):615–627.

50. Sabbagh JJ, Kinney JW, & Cummings JL (2013) Animal systems in the development of treatments for Alzheimer’s disease: challenges, methods, and implications. Neurobiology of aging 34(1):169–183.

51. Miyakawa T (2020) No raw data, no science: another possible source of the reproducibility crisis. Molecular brain 13(1):24.

52. Vrselja Z, et al. (2019) Restoration of brain circulation and cellular functions hours post-mortem. Nature 568(7752):336–343.

53. Choe MS, et al. (2021) A simple method to improve the quality and yield of human pluripotent stem cell-derived cerebral organoids. Heliyon 7(6):e07350.

54. Qian X, et al. (2020) Sliced Human Cortical Organoids for Modeling Distinct Cortical Layer Formation. Cell stem cell 26(5):766–781 e769.

55. Iyer A, Rajkumar V, Sadasivan D, Bruce J, & Gilfillan I (2007) No one is dead until warm and dead. The Journal of thoracic and cardiovascular surgery 134(4):1042–1043.

56. Shen KF & Schwartzkroin PA (1988) Effects of temperature alterations on population and cellular activities in hippocampal slices from mature and immature rabbit. Brain research 475(2):305–316.

57. Graham BA, Brichta AM, & Callister RJ (2008) Recording temperature affects the excitability of mouse superficial dorsal horn neurons, in vitro. Journal of neurophysiology 99(5):2048–2059.

58. Pasikova NV, Mednikova Iu S, & Averina IV (2010) [The temperature regulation of the spontaneous activity frequency and the concomitant changes of the cortical neurons’ spike amplitude]. Rossiiskii fiziologicheskii zhurnal imeni I.M. Sechenova 96(3):315–324.

59. Proskurina EY & Zaitsev AV (2021) Photostimulation activates fast-spiking interneurons and pyramidal cells in the entorhinal cortex of Thy1-ChR2-YFP line 18 mice. Biochemical and biophysical research communications 580:87–92.

60. Gunaydin LA, et al. (2014) Natural neural projection dynamics underlying social behavior. Cell 157(7):1535–1551.

61. Moore JJ, et al. (2017) Dynamics of cortical dendritic membrane potential and spikes in freely behaving rats. Science 355(6331).

62. Hai A, Shappir J, & Spira ME (2010) Long-term, multisite, parallel, in-cell recording and stimulation by an array of extracellular microelectrodes. Journal of neurophysiology 104(1):559–568.

63. Dipalo M, et al. (2017) Intracellular and Extracellular Recording of Spontaneous Action Potentials in Mammalian Neurons and Cardiac Cells with 3D Plasmonic Nanoelectrodes. Nano letters 17(6):3932–3939.

