## Supplementary figures and images for "Investigation of neural functional connectivity in thick acute mouse brain slices with novel multi-region 3D neural probe arrays"

### Supplemental 1

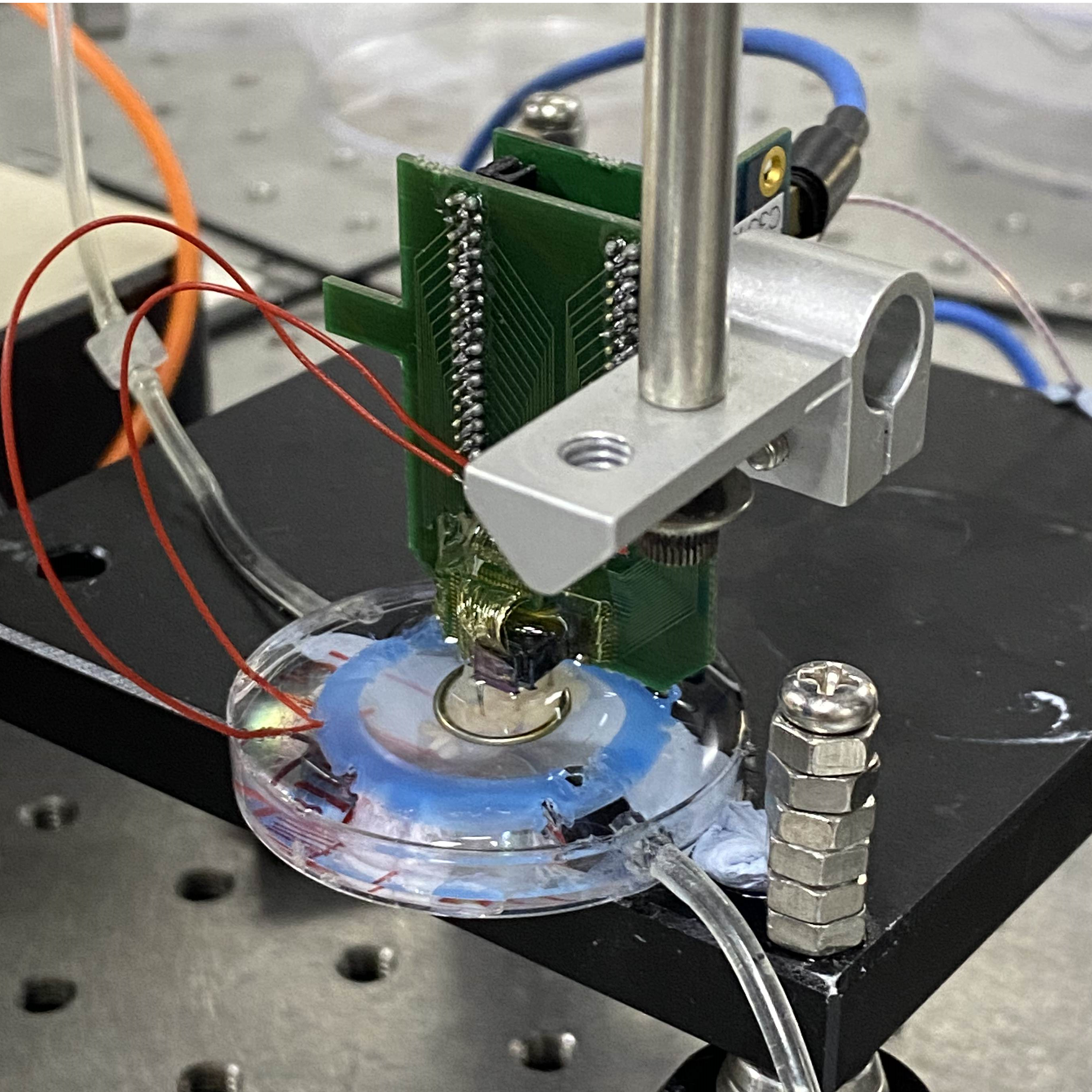

### Supplemental 2

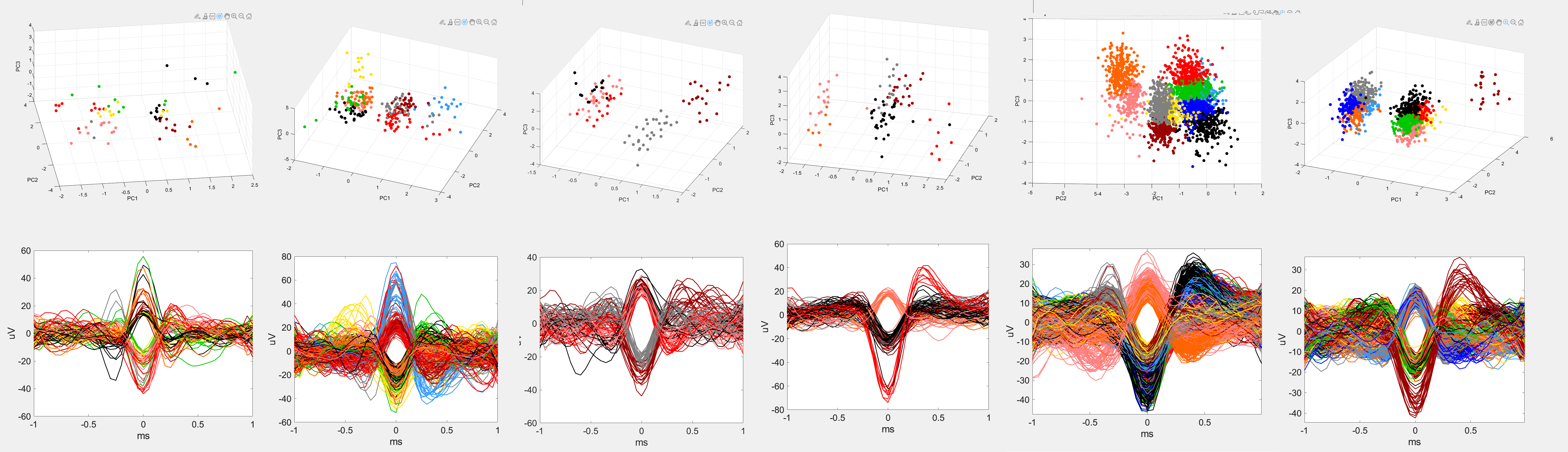

### Supplemental 3

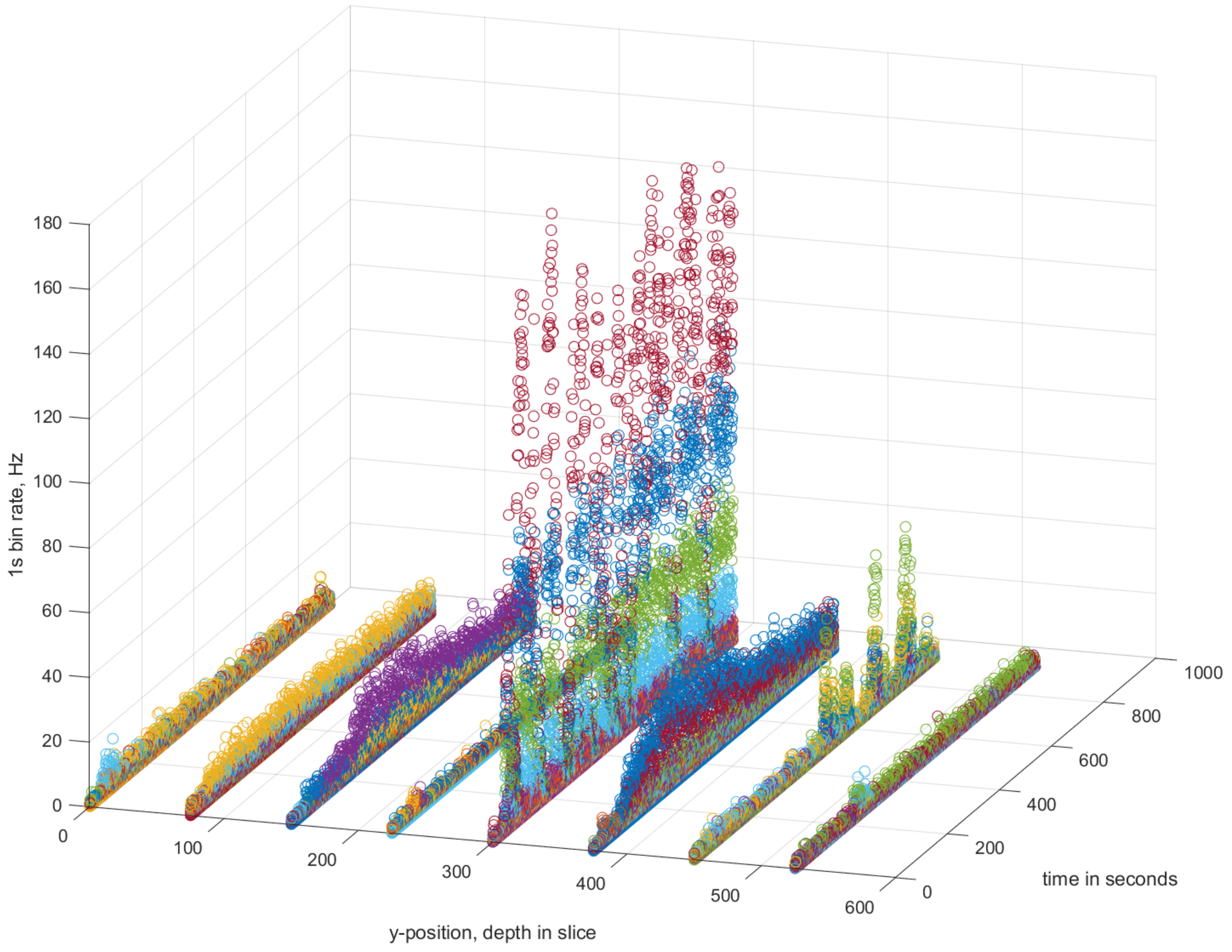

### Supplemental 4

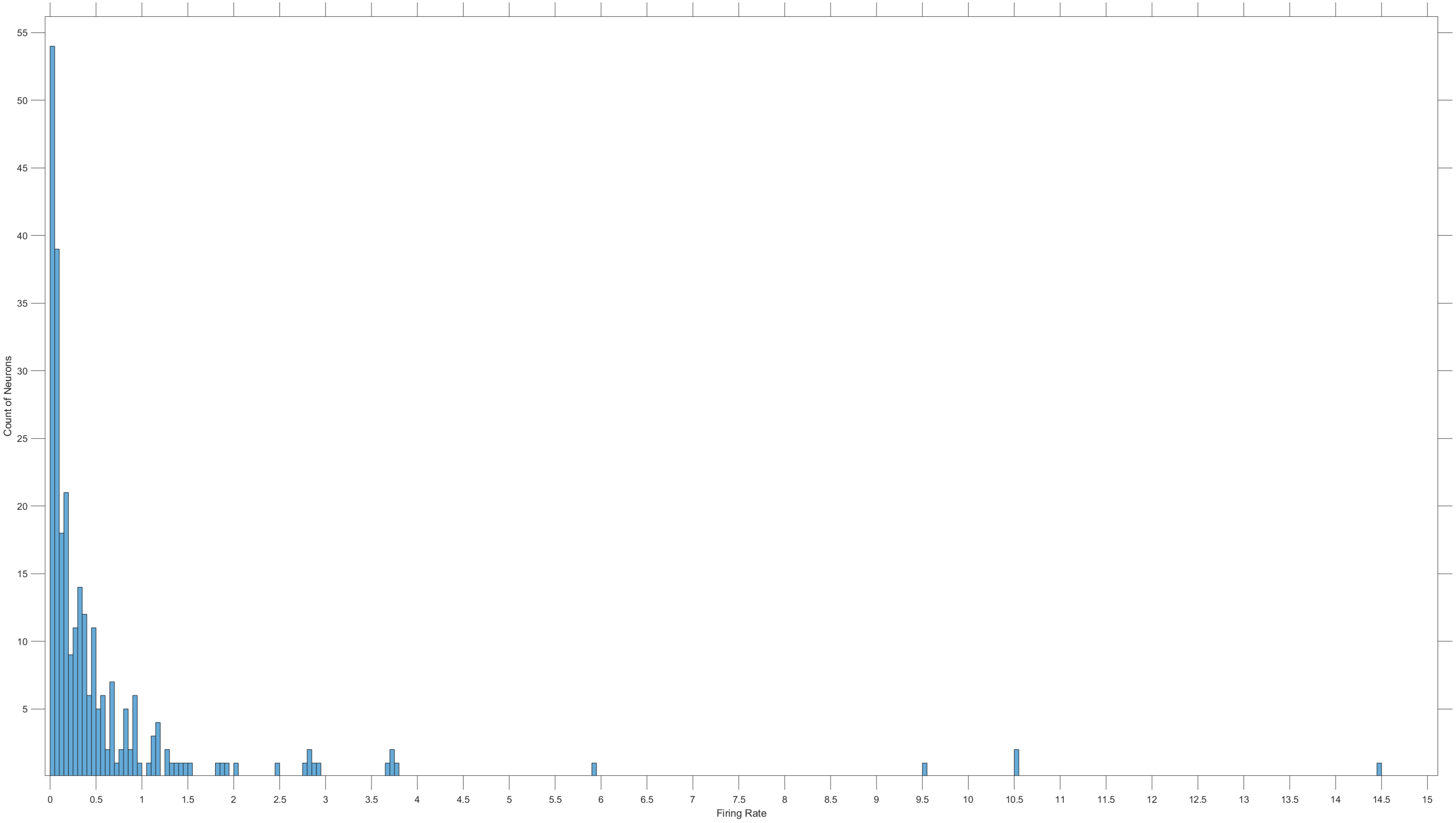

### Supplemental 5

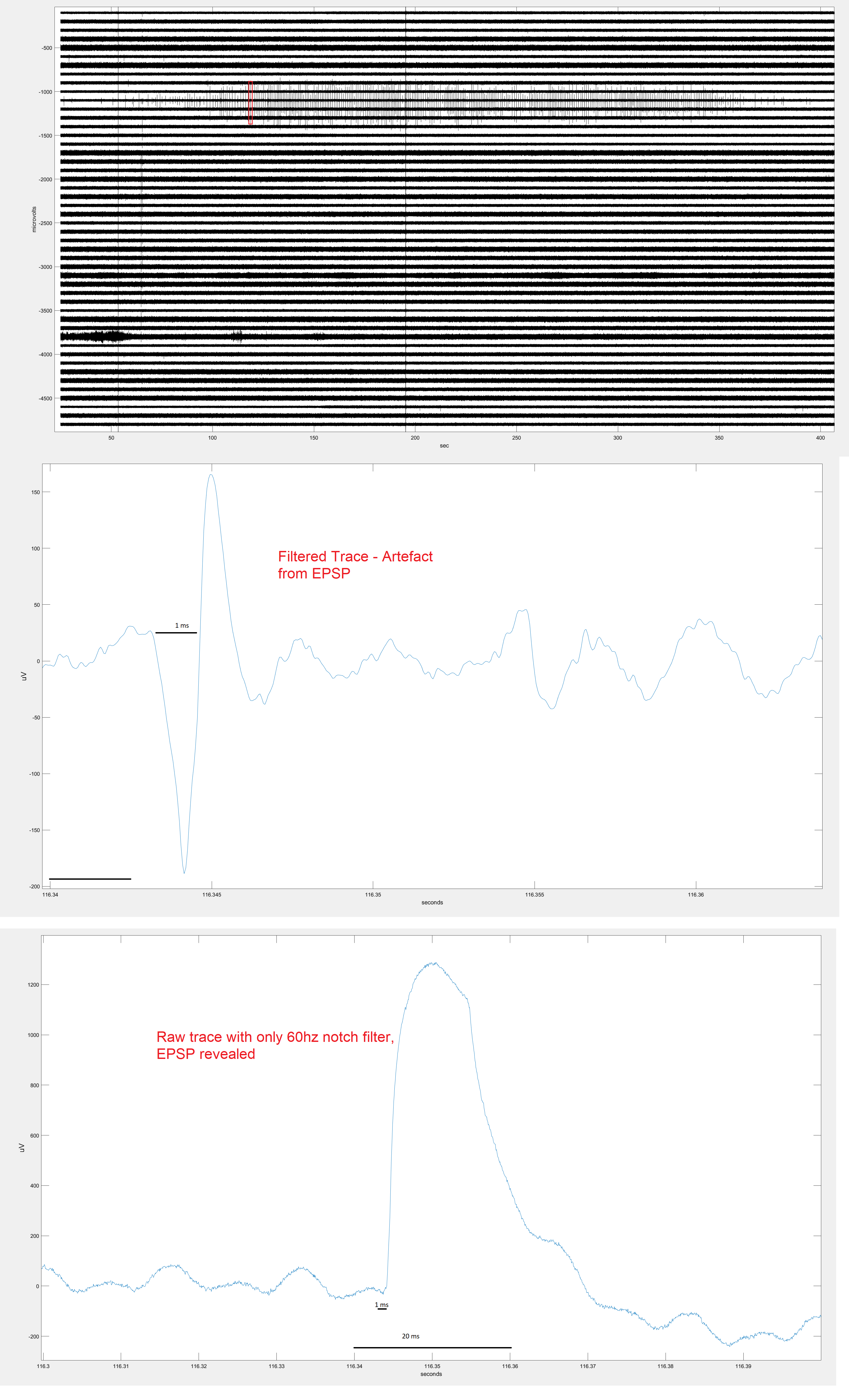
